# Peptide nucleic acid (PNA) and DNA hybrid three-way junctions and mesojunctions

**DOI:** 10.1101/2025.05.15.653195

**Authors:** Bharath Raj Madhanagopal, Akul Patel, Hannah Talbot, John A. Geary, Krishna N. Ganesh, Arun Richard Chandrasekaran

## Abstract

Three-way junctions are simple and fundamental structural motifs that impart the typical branching property in most DNA nanostructures. While conventional three-way DNA junctions are well-known, mesojunctions are relatively unexplored. Here, we report the synthesis of peptide nucleic acid (PNA)/DNA hybrid three-way conventional and mesojunctions, containing a 14 bp duplex DNA arm and two PNA/DNA hybrid arms. We show that the PNA/DNA mesojunction can be assembled in magnesium-free buffers containing low concentrations of calcium or sodium salts. PNA/DNA hybrid junctions and mesojunctions showed higher thermal stability compared to the DNA versions. Further, PNA/DNA hybrid junctions assembled in sodium exhibited higher nuclease resistance against DNase I. Our results pave the way for PNA/DNA hybrid three-way junctions to be integrated into complex DNA nanostructure designs to improve their structural and enzymatic stability.

---

Sequence-based self-assembly of DNA has led to the construction of various nanostructures including polyhedra, nanowires, and periodic lattices.^1,2^ A defining feature of DNA nanostructures is the occurrence of DNA crossovers and branch points that allow the construction of multidimensional objects and arrays, with most branch points containing a three-way junction or a four-way junction.^3^ The conventional three-way junction is one of the fundamental structural motifs used routinely in the design of complex nucleic acid nanostructures.^4^ It has three double-helical arms emanating from the center of the structure. When visualized as the convergence of the double-helices into a central triangle, the helix axes of all three double-helical arms, one at each vertex, pass through the center of the triangle (**Figure 1a**). Thus, a conventional junction is denoted as a 3_3_ junction, where the number corresponds to the number of strands in the complex, and the subscript indicates the number of radial axes (**Figure 1b**).^5^ In a mesojunction, the axis of only one of the three arms passes through the center, and the helix axes of the other two arms are circumferential (a 3_1_ junction) (**Figure 1c**). In addition to three-way junctions, four-way DNA antijunctions and mesojunctions have also been reported.5,6 While several studies have explored various aspects of the canonical DNA three-way and four-way junctions,^7^ including their three-dimensional structure,^8,9^ thermal and thermodynamic stability,^10^ and enzymatic stability,^11,12^ there are relatively fewer studies on mesojunctions. Some studies have used variations of DNA mesojunctions in the construction and reconfiguration of DNA nanostructures.^13^ Assembly of mesojunctions using artificial DNA analogs such as peptide nucleic acids (PNA) can be useful to improve the assembly and stability of nanostructures based on this motif and provide additional functionality for biological applications.^14^

**Figure 1.**
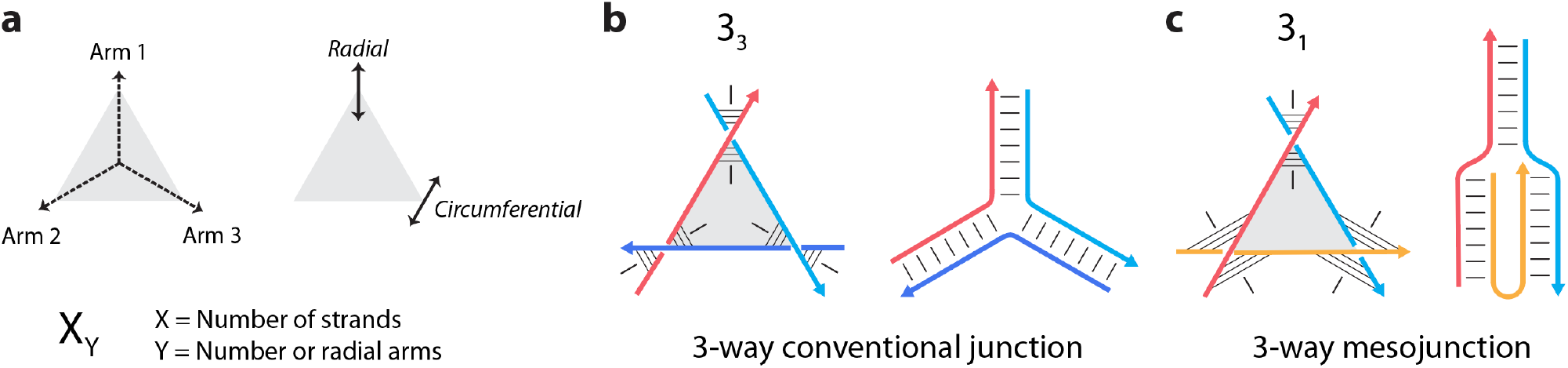
DNA conventional junctions and mesojunctions. (a) Schematic showing the three arms of the 3-way junction emanating from the center. The arms of the junction can point radially or be circumferential to the vertex of the triangle. The standard nomenclature for the junctions is a number indicating the total number of strands followed by a subscript for the number of radial arms. (b) A canonical three-way junction has all three arms passing through the center (a 3_3_ junction). (c) A three-way mesojunction has only one radial arm out of the three arms (a 3_1_ junction). Arrowheads indicate the 3’ ends of strands. The thin black lines indicate the base pairing between the antiparallel strands, and the lines perpendicular to the base pairs indicate the helical axis.

PNA is a DNA analog with an achiral backbone consisting of uncharged repeats of *N*-(2-aminoethyl) glycine units linked by amide bonds (**Figure 2a**).^15,16^ PNA binds to DNA strands with high affinity, shows high sequence fidelity (**Figure 2b**), has low dependency on ionic strength,exhibits high chemical stability, and displays resistance to both nucleases and proteases.^17^ In prior work, PNA has been incorporated into DNA nanostructures such as a four-way junction,^18^ a double crossover DNA motif,^19,20^ a triple crossover DNA motif,^21^ a tetrahedron,^22^ and a rectangular DNA origami.^21^ In most of these cases, the PNA strand was attached to single-stranded extensions protruding out of the structure rather than directly incorporated as part of the structure. These studies have looked at the structural changes induced by PNA on an otherwise DNA-based nanostructure (such as changes in helicity^19^) and use PNA as a tether to attach cargoes on DNA nanostructures (such as a PNA-peptide conjugate^22^). Structures such as a three-way junction and helical bundles have also been made entirely out of PNA.^23,24^ However, the junctions studied in these cases are conventional junctions, and PNA/DNA hybrid mesojunctions have not been investigated. In this work, we constructed PNA/DNA hybrid conventional three-way junctions and mesojunctions and compared their assembly and characteristics to all-DNA junctions. We further show the feasibility of assembling PNA/DNA hybrid junctions and mesojunctions in magnesium-free conditions by substituting magnesium with calcium or sodium ions in the buffer. PNA/DNA hybrid junctions assembled in a sodium-containing buffer also show enhanced nuclease resistance compared to junctions assembled in magnesium-containing buffer.

**Figure 2.**
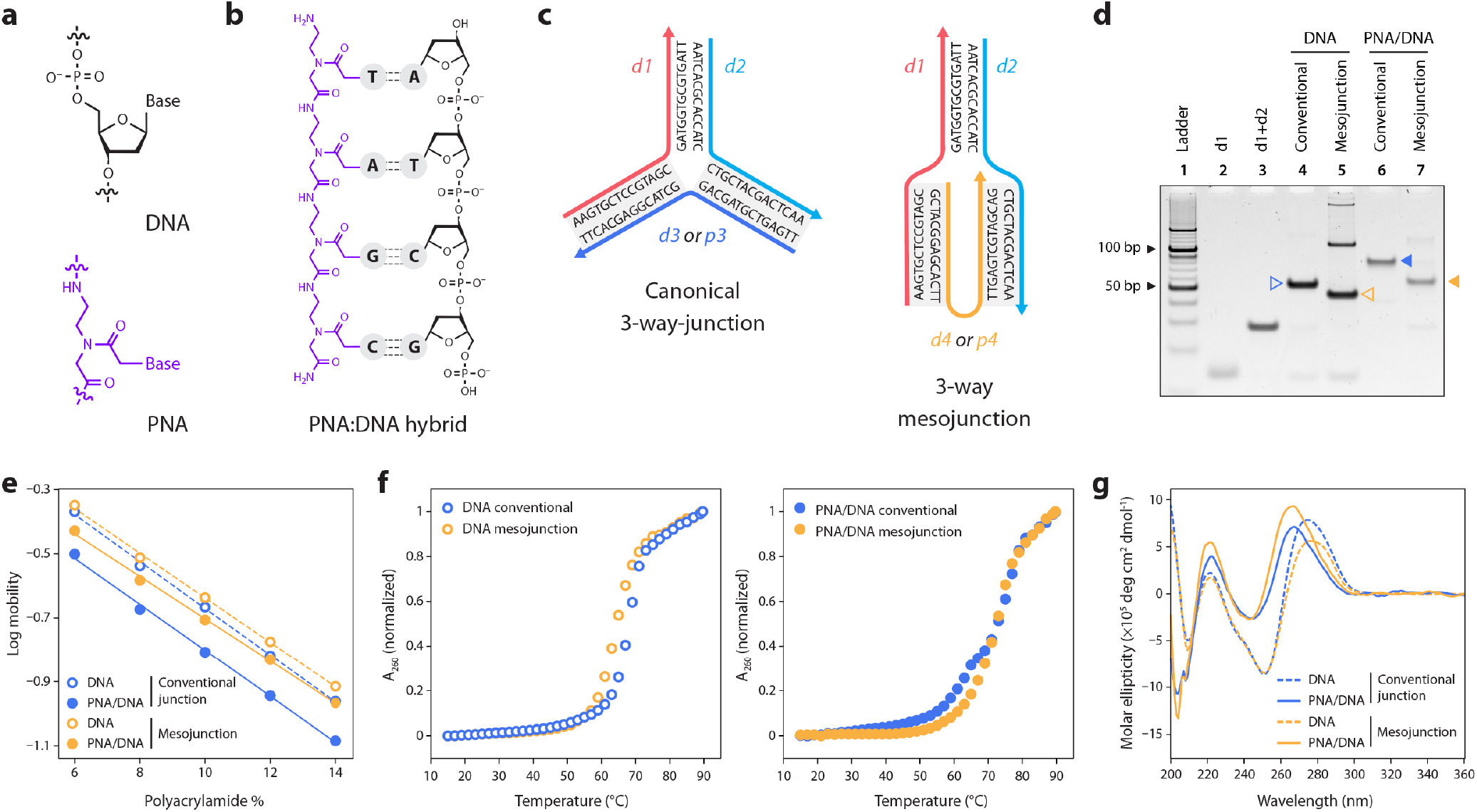
Assembly of PNA/DNA hybrid junctions. (a) Structure of DNA and peptide nucleic acid (PNA). (b) Base pairing between PNA and DNA strands. (c) Design and sequences of the three-way conventional junction and mesojunction. (d) Gel showing the assembly of the junctions. The mesojunctions for both DNA and PNA/DNA migrate faster than the conventional junctions. PNA/DNA hybrid junctions in general have reduced mobility compared to the DNA versions. (e) Ferguson plot for diKerent junctions. (f) Thermal melting curves of conventional and mesojunctions. (g) CD spectra of conventional and mesojunctions.

We first designed DNA three-way junctions with different arm lengths based on previously reported sequences^5^ (**Figure S1, Table S1**). The respective conventional junctions and mesojunctions have the same base composition but differ by the sequence of only one component strand, which determines the number of radial axes in the junctions (**Figure 2c** and **Figure S1**). In the design shown in Figure 2c, strands d1 and d2 are common for both structures, with strand d3 completing the conventional junction and strand d4 completing the mesojunction. We constructed the conventional junction and mesojunction with 12 bp, 14 bp, and 16 bp arms in tris-acetate-EDTA (TAE) buffer with 12.5 mM magnesium acetate and validated assembly using non-denaturing polyacrylamide gel electrophoresis (PAGE). Assembly yields of the conventional junctions were relatively the same regardless of the duplex arm lengths, whereas the assembly yield of the mesojunction with 14 bp arm was the highest (**Figure S2**). Further, the assembly yields of DNA mesojunction were relatively lower than the conventional junction, and we observed the formation of higher-order structures, a result that is consistent with previously reported DNA mesojunctions.^5^ Based on the assembly yields, we chose the junction design with 14 bp arms to construct the PNA/DNA hybrid junctions.

To construct the PNA/DNA hybrid junctions, we replaced one of the DNA strands in the junction to be a PNA (p3 instead of d3 for the conventional junction and p4 instead of d4 for the mesojunction) (**Figure 2c**). We assembled the PNA/DNA hybrid junctions using a protocol similar to the one we used for the DNA junctions and confirmed assembly using non-denaturing PAGE (**Figure 2d**). We confirmed the formation of the junctions by comparing their mobility to complexes formed from subsets of the component strands (**Figure S3**). The PNA/DNA hybrid conventional junction (Figure 2d, lane 6) migrated slowly in the gel compared to the DNA junction (lane 4), perhaps due to the neutral backbone of the PNA. Similarly, the PNA/DNA mesojunction (lane 7) migrated slowly compared to the DNA mesojunction (lane 5). We note that the positively charged single-stranded PNA do not run on the gel and the complexes with only PNA/DNA hybrid duplex arms are not stained by GelRed, the stain used to visualize the bands. PNA/DNA hybrids with one double stranded DNA duplex arm are stained weakly in comparison to all-DNA structures (**Figure S3**). We then analyzed the junctions using a Ferguson plot, where the log (mobility) as a function of polyacrylamide concentration provides information about the charge on the molecule and its molecular weight (**Figure 2e** and **Figure S4**).^25^ The slope of the DNA mesojunction differs from that of the DNA conventional junction (**Table S2**), consistent with previous analysis of DNA three-way junctions.^5^ Similarly, the conventional and mesojunctions of PNA/DNA hybrids exhibited different slopes, indicating that the different orientations of the strands (and thus the helical arms) resulted in different surface areas. The Y-intercepts of the curves on a Ferguson plot correspond to the charge on the molecule. As expected, the Y-intercepts of PNA/DNA hybrid junctions varied from those of DNA junctions. While there was no major difference between the Y-intercepts of the conventional and mesojunctions of DNA, the values were different for the conventional and mesojunctions of the PNA/DNA hybrids. This suggests that counterions might play a distinct role in the structure of the conventional and mesojunctions of PNA/DNA hybrids.

After confirming the assembly of the conventional junctions and mesojunctions, we measured their thermal stability by monitoring the temperature-dependent changes in the absorbance at 260 nm (**Figure 2f**). In the case of DNA structures, the conventional junction was slightly more stable (T_m_ = 67.8 °C) than the mesojunction (T_m_ = 64.0 °C), consistent with a previous study^5^ (**Table S3**). In comparison to the DNA junction, the PNA/DNA hybrid junctions had a higher melting temperature (76 °C for PNA/DNA conventional junction and 72.9 °C for PNA/DNA mesojunction). As observed in the previous study,^5^ the thermal curves of the DNA junctions indicated premelting transitions, with PNA/DNA conventional junction showing a prominent transition centered around 64 °C. Similar melting profiles have been observed for bimodal PNAs earlier.^26^ The observation of the low-melting transition indicates that the dissociation of the PNA/DNA hybrid conventional junction is not a two-state phenomenon, with perhaps the DNA arm melting at a lower temperature and the PNA/DNA hybrids melting at a higher temperature. Interestingly, the PNA/DNA mesojunction does not show a distinct low-melting transition. However, it is not clear if this is due to the presence of higher-order structures in the assembly. Circular dichroism spectra of the DNA junctions were similar and reflected the underlying B-form DNA duplex arms (**Figure 2g**). Compared to the DNA junctions, the CD spectra of the PNA/DNA junctions showed a blue-shift in the positive band from ∼275 nm to ∼265 nm and negative band from 250 nm to ∼240 nm, which is typically observed in PNA/DNA hybrids.^27^

We then studied the effect of counter ions on the assembly of the PNA/DNA hybrid junctions. The assembly of the DNA mesojunction required at least 5 mM Mg^2+^ concentration, with no major improvement at higher Mg^2+^ concentrations (**Figure 3a**). For the PNA/DNA mesojunction, we observed a slight decrease in the intensity of the bands with increasing concentrations of Mg^2+^. We then tested the assembly of the junctions in other metal ions. Our recent work showed that DNA motifs can be assembled through an annealing process or isothermally in a wide variety of cations instead of the typically used Mg^2+^.^28,29^ Here, we assembled the all-DNA and PNA/DNA hybrid conventional junctions and mesojunctions in Ca^2+^ and Na^+^ in addition to the assembly in Mg^2+^. The assembly yields of all the junctions in 10 mM Ca^2+^ were similar to the assembly yields in 10 mM Mg^2+^ (**Figure 3b-c**), indicating that PNA/DNA hybrid structures can be assembled in Ca^2+^-containing buffers. In Na^+^, the DNA and PNA/DNA hybrid conventional junctions were well formed, with yields similar to those assembled in Mg^2+^. The assembly yields of the conventional junctions were also similar in 10 and 100 mM Na^+^. For the DNA mesojunction, the assembly yield was poor in the buffer containing 10 mM Na^+^, with even lower yields in 100 mM Na^+^ (**Figure 3b**, lanes 8 and 9). The monovalent Na^+^ ion appears to favor the formation of higher-order structures rather than the folding of the strands within the structure, which is required to form isolated junctions. This observation highlights the need for divalent ions to form stable DNA mesojunctions, where the axes of two arms of the junction are brought closer by the bridging strand. Perhaps, the divalent ions are needed to screen the negative charges between the backbones of the juxtaposed duplexes and/or to overcome kinetic traps during the assembly. In contrast, the PNA/DNA hybrid mesojunction formed with higher assembly yields in buffer containing Na^+^ (**Figure 3c**, lanes 8 and 9). Unlike DNA mesojunctions, stable PNA/DNA mesojunctions can be formed at 10 mM or 100 mM Na^+^ concentrations.

**Figure 3.**
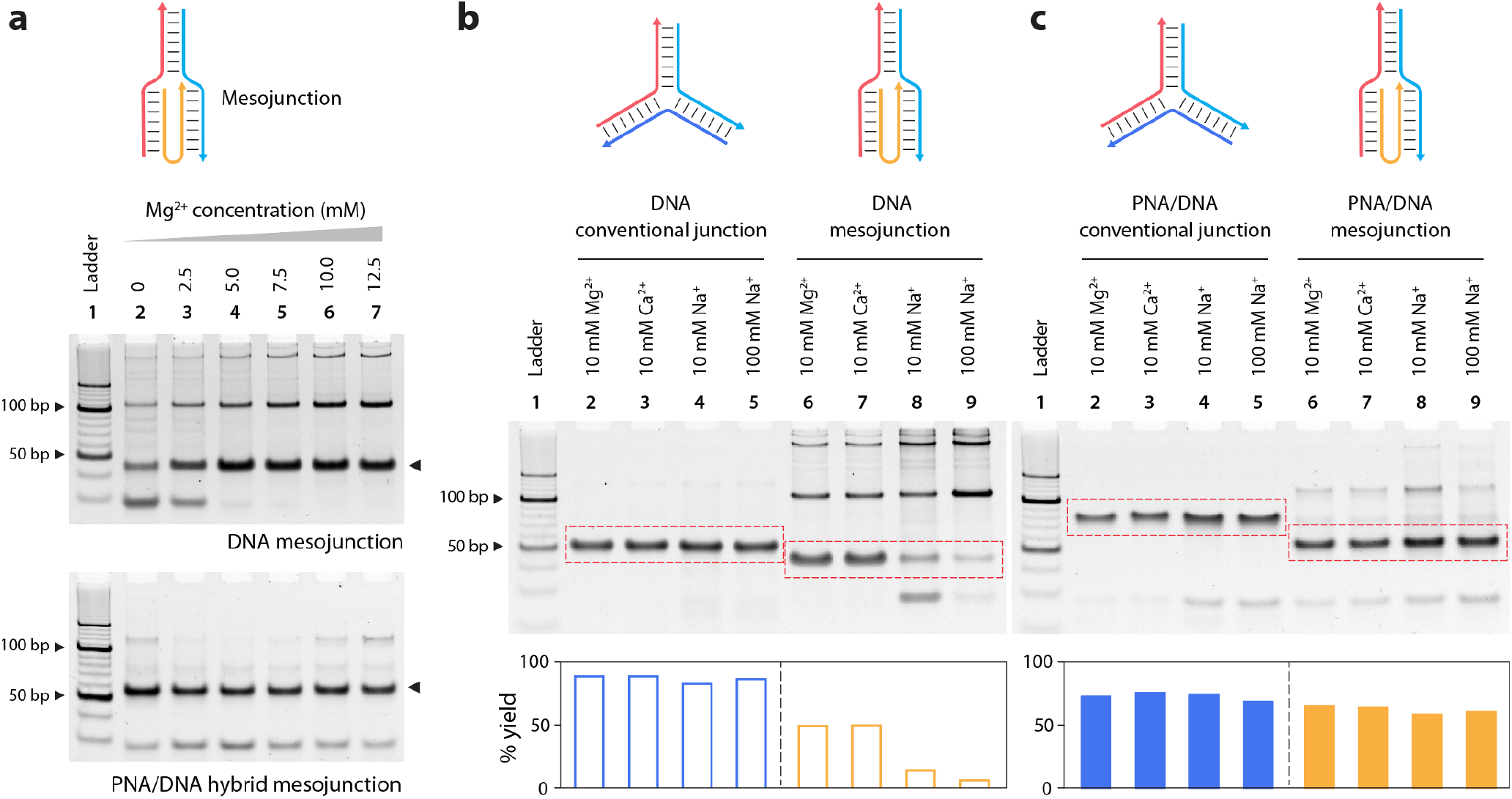
Effect of counter ions on junction assembly. (a) Effect of Mg^2+^ concentration on the assembly of DNA and PNA/DNA mesojunctions. (b) Assembly of DNA and (c) PNA/DNA conventional and mesojunction using Mg^2+^ (10 mM), Ca^2+^ (10 mM) and Na^+^ (10 and 100 mM).

Since PNA has a synthetic backbone, it is not recognized by nucleolytic enzymes and nanostructures made using PNA are likely to show higher biostability against the common nucleases.^30^ We tested whether the PNA/DNA hybrid junctions are able to resist nucleolytic attack by DNase I. First, we studied the junctions assembled in TAE buffer with 12.5 mM Mg^2+^ by incubating them with different amounts of DNase I and measuring the intact fraction of the band corresponding to the structures on a non-denaturing gel. The PNA/DNA hybrid junctions showed slightly higher nuclease resistance than the DNA junctions (**Figure 4** and **Figure S5**). In our prior work, we showed that DNA motifs assembled in monovalent ions are more biostable compared to those assembled in divalent ions.^28^ To verify this phenomenon in PNA/DNA hybrid junctions, we tested the nuclease resistance of junctions assembled in 100 mM Na^+^. We observed that the PNA/DNA junctions assembled in Na^+^ were much more nuclease resistant (88% and 64% intact for conventional and mesojunction, respectively, against 0.2 U/µl DNase I) compared to the junctions assembled in Mg^2+^ (13-32% intact).

**Figure 4.**
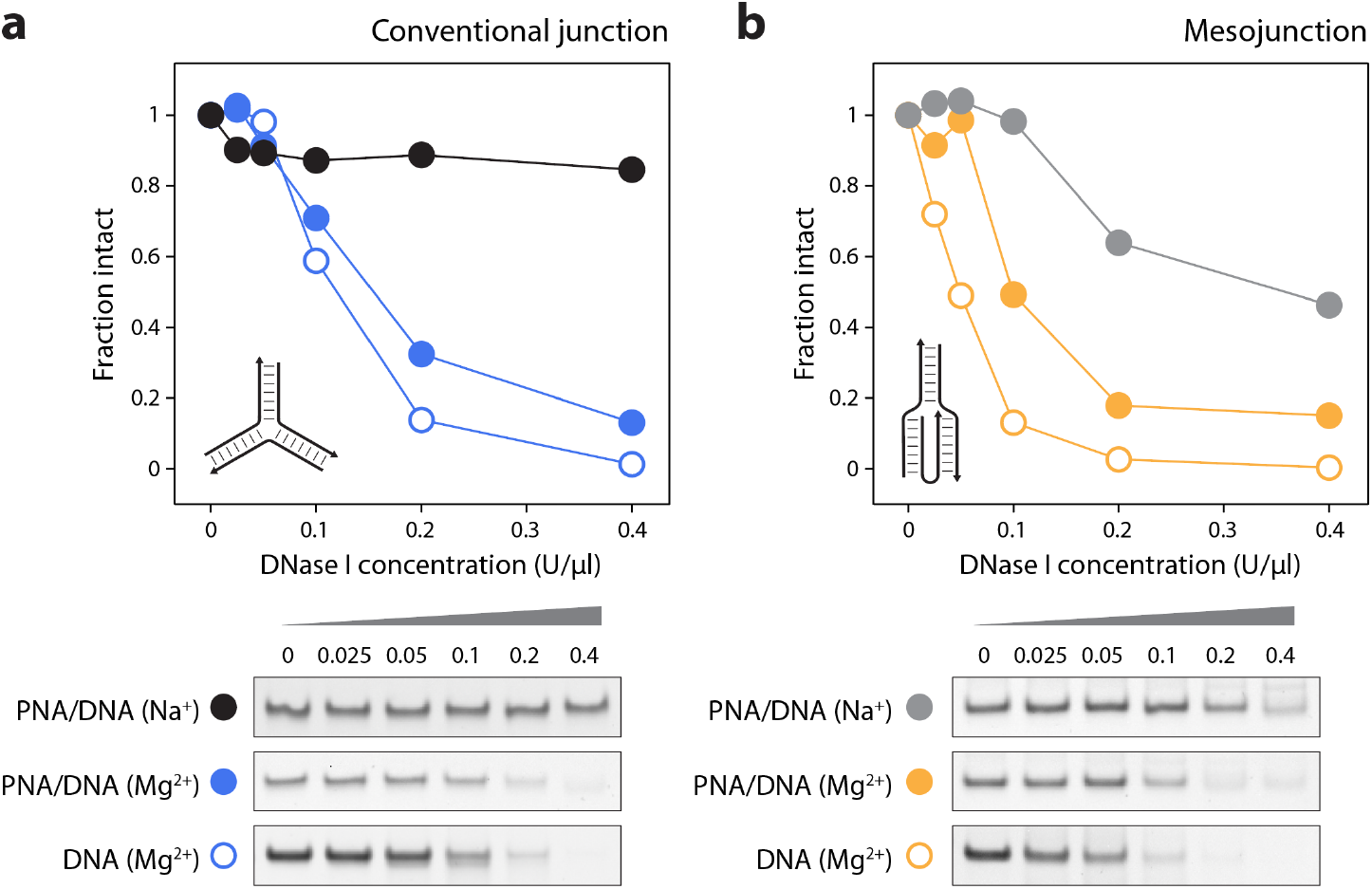
Nuclease resistance of PNA/DNA junctions. Degradation profiles of DNA junctions assembled in Mg^2+^, PNA/DNA junction assembled in Mg^2+^, and PNA/DNA junction assembled in Na^+^. (a) shows degradation profiles against DNase I for conventional junctions and (b) shows degradation profiles against DNase I for mesojunctions.

Synthetic DNA mimics such as peptide nucleic acids, locked nucleic acids, threoninol nucleic acid, and threose nucleic acids offer several advantages, including increased thermal stability and biostability.^31–34^ The use of these structural analogs to construct DNA nanostructures can expand the scope of these functional materials in various applications. Our results show that it is possible to assemble PNA/DNA hybrid junctions and mesojunctions with good assembly yields by using PNA as one of the strands. Assembly using PNA reduces the reliance on high divalent concentrations for the assembly, specifically for PNA/DNA mesojunctions that assemble with high yields in Na^+^-containing buffer. In contrast, Na^+^ precludes the assembly of all-DNA mesojunction. We also showed that PNA/DNA hybrid structures assembled in Na^+^ are more biostable. This information is useful for creating PNA-based nucleic acid nanostructures for biological applications where the protective effect of Na^+^ for mesojunction-containing structures is an advantage over all-DNA structures.

The neutral backbone of PNA allows us to harness its DNA-like programmable self-assembling property in non-aqueous solvents, extending the application value of DNA nanotechnology.^24^ In addition to the previously shown interesting self-assembling properties of di-PNAs,^35^ bimodal PNAs,^36^ and branched PNAs,^37^ the present work highlights the versatile range of self-assembled structures that can be designed with PNA. A major limitation of using PNA is the tendency of the molecules to aggregate, especially when the length of the PNA is as long as the 28-mer strands used in this work. Here, we introduced two L-lysines at the C-terminus to improve the aqueous solubility of the molecule, yet we observed that the assembly yields of PNA/DNA hybrid junctions were inconsistent. Modified PNAs with higher aqueous solubility might be more useful for constructing nanostructures.^17^ To incorporate artificial nucleic acids in the design of DNA nanostructures, it is important to understand the secondary structures adopted by these modified nucleic acids and nucleic acid mimics.^19^ In open-ended structures such as a three-way junction, the geometry of the non-B-DNA double helix formed by PNA and DNA is not restrictive, allowing us to study the properties of such hybrid nanostructures. For future implementation of these hybrid designs in larger nanostructures, the helical parameters of the hybrid double helix may guide the position and integration of the hybrid duplex segments in the design of nucleic acid nanostructures. Intricate nanostructure design using PNA analogs may also result in robust nanostructures^38^ with better in vivo activity,^39^ allowing the use of PNA-based architectures in nucleic acid nanostructure design for biological applications.

## Competing interests

The authors have no competing interests.

## Acknowledgments

Research reported in this publication was supported by the National Institutes of Health (NIH) through National Institute of General Medical Sciences (NIGMS) under award number R35GM150672 to A.R.C.

## Supporting information

Experimental procedures, additional results and DNA sequences used.

## Notes

### Competing Interest Statement

The authors have declared no competing interest.

