## Supporting information for "Peptide nucleic acid (PNA) and DNA hybrid three-way junctions and mesojunctions"

### Methods

#### Preparation of DNA and PNA junctions

DNA strands were purchased from Integrated DNA Technologies (IDT) with standard desalting and used without further purification. Stock solutions of DNA oligonucleotides were prepared in nuclease-free water, and their concentrations were estimated using Nanodrop 2000 spectrophotometer (Thermo Scientific) by measuring their absorbance at 260 nm. Peptide nucleic acid (PNA) strands were purchased from Bio-Synthesis (Lewisville, TX). The PNA strands were purified by RP-HPLC (>90%) and characterized by MALDI-TOF by the manufacturer. The metal salts used in the buffers were calcium chloride dihydrate (>99%, Sigma), magnesium acetate tetrahydrate (>99%, Sigma), magnesium chloride hexahydrate (>99%, VWR), and sodium chloride (>99%, Millipore).

The DNA junctions and PNA/DNA junctions were assembled by mixing the component strands in the specified molar concentrations in tris-acetate EDTA buffer containing 40 mM Tris base (pH 8.0), 20 mM acetic acid, 2 mM EDTA, and 12.5 mM magnesium acetate (1× TAE-$Mg^{2+}$). To test the junction assembly in different salts, the equimolar mixtures of the component strands were prepared in 1× TAE buffer with the required amounts of magnesium chloride, calcium chloride, or sodium chloride. The solutions were annealed in a thermal cycler with the following steps: 95°C for 3 minutes, 65°C for 20 minutes, 45°C for 20 minutes, 37°C for 30 minutes, 20°C for 30 minutes, and 4°C. The solutions were stored at 4°C.

#### Non-denaturing polyacrylamide gel electrophoresis

Polyacrylamide gels were prepared using 19:1 acrylamide solution (National Diagnostics) in 1× TAE-$Mg^{2+}$ buffer. The annealed samples were mixed with 10× loading dye containing bromophenol blue and glycerol before being loaded on the gel. The gels were run at 4 °C with 1× TAE-$Mg^{2+}$ as the running buffer at 100 V for 90 min to 2 hours, as required. The gels were stained with 0.5× GelRed (Biotium) in water for 20 min in dark, destained in water for 10 min, and imaged on a Bio-Rad Gel Doc XR+ imager using the default settings for GelRed with UV illumination. The intensities of the bands and their relative migration on the gels were measured using ImageLab software (BioRad).

#### Circular dichroism (CD) spectroscopy

DNA hybrid junctions were prepared in 1× TAE-$Mg^{2+}$ buffer at a concentration of 10 μM. PNA/DNA junctions were prepared in the same buffer at a concentration of 3 μM. 200 μl of the junctions were taken in a 1 mm thick quartz cuvette, and the CD spectra were recorded on a Jasco J-815 CD spectrometer with a scan speed of 100 nm/min. The bandwidth and digital integration time were set at 1 nm and 1 s, respectively. The average of three accumulations was used for further analysis. The spectra were smoothened using with Savitzky-Golay method, and molar ellipticity was calculated for all the samples using the Spectra Manager software (JASCO).

#### Thermal melting studies

UV-thermal melting of the junctions was performed on a Cary 3500 UV-Visible spectrophotometer (Agilent). The DNA junctions were annealed at a concentration of 2.5 μM, and the PNA/DNA hybrid junctions were prepared at a concentration of 1.5 μM in 1× TAE-$Mg^{2+}$ buffer containing 12.5 mM magnesium acetate. The solutions were heated from 15 °C to 90 °C at a rate of 0.5 °C/min, and the absorbance at 260 nm was recorded. The melting curves were normalized. Boltzmann function did not produce good fits for some of the samples, and Double Boltzmann function using OriginPro. The maximum points on the first derivative curves were noted as the melting temperature.

**Enzyme degradation studies**

The DNA and PNA/DNA hybrid junctions were prepared in 1× TAE containing 10 mM $MgCl_2$, 10 mM NaCl, or 100 mM NaCl. 9 µl of the assembled junction samples were mixed with DNase I reaction buffer provided by the vendor (1× final). 9 µl of this mixture was added to 1 µl of 0.025 to 0.4 U/µl DNase I (New England BioLabs) and incubated at 37 °C for 30 min. The samples were loaded on non-denaturing PAGE gels and run for 2 hours.

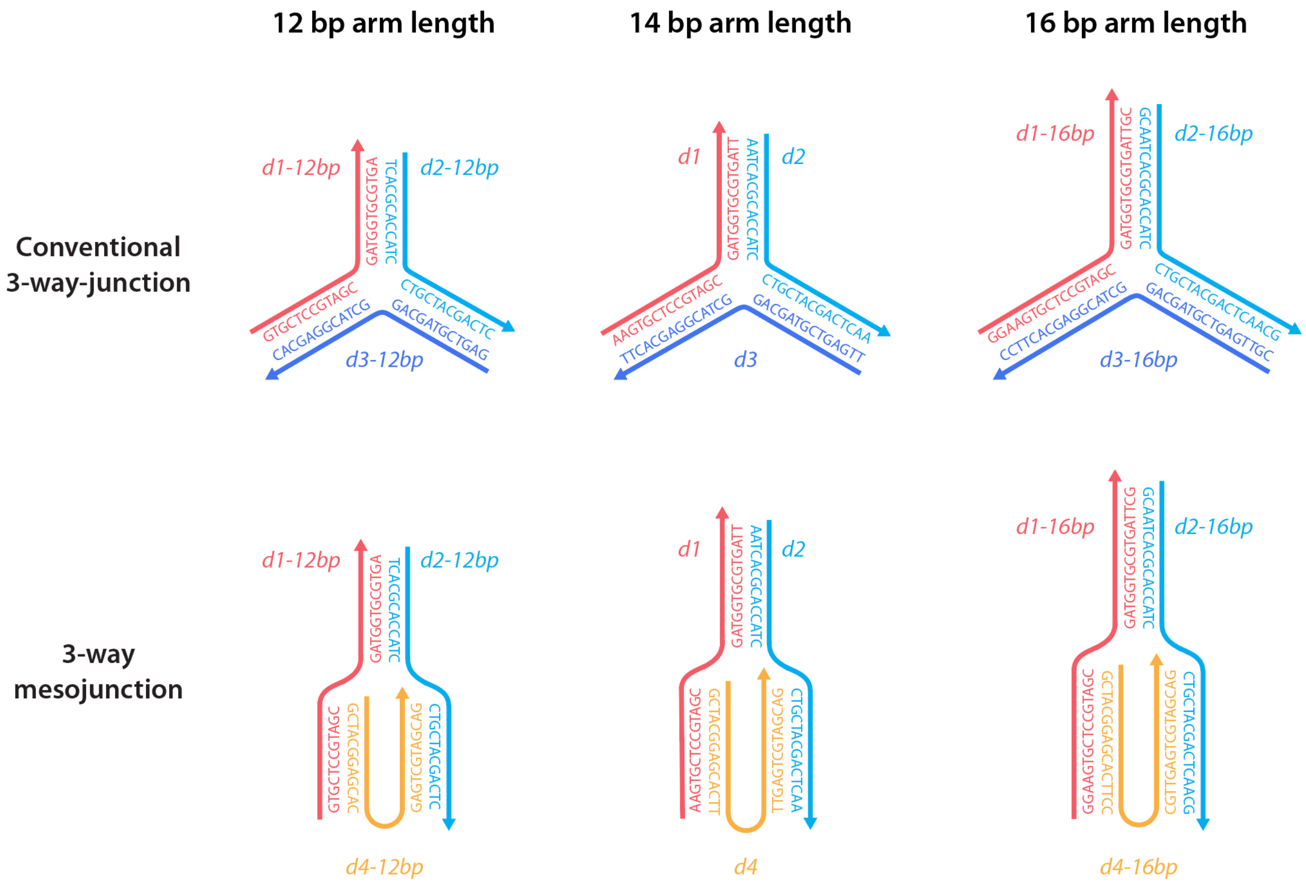


**Figure S1**. Design of the junctions and their sequences.

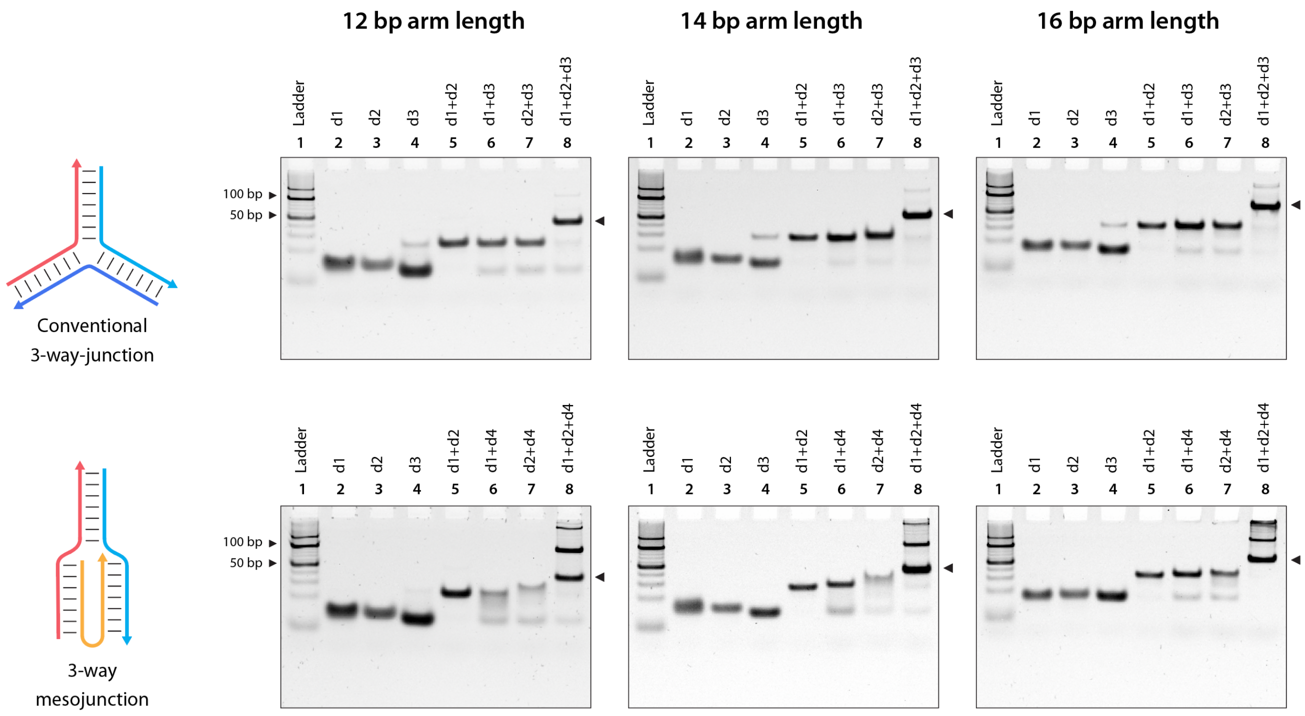


**Figure S2**. Non-denaturing gels showing the assembly of DNA conventional junctions and mesojunctions with different arm lengths.

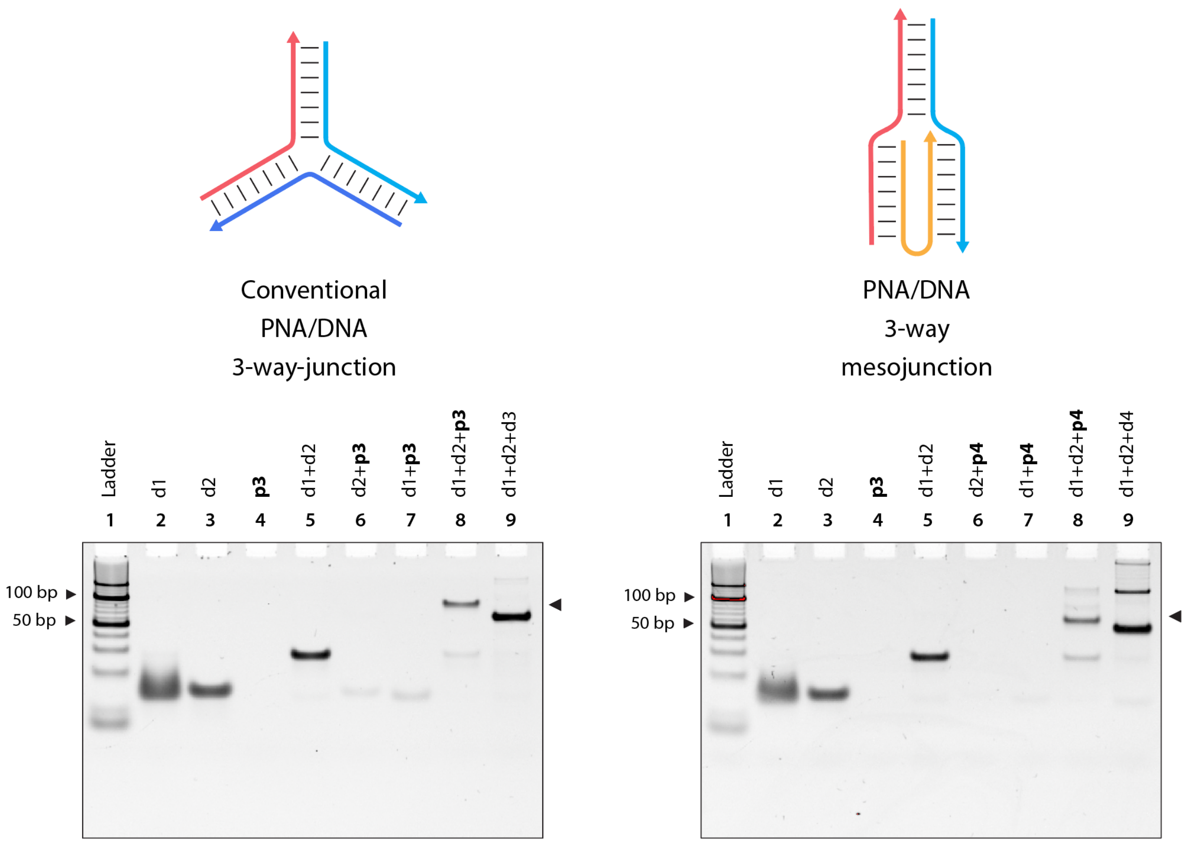


**Figure S3**. Assembly of PNA/DNA hybrid conventional junction and mesojunction. Strand combinations with **d** denote DNA strands, and **p** denote PNA strands.

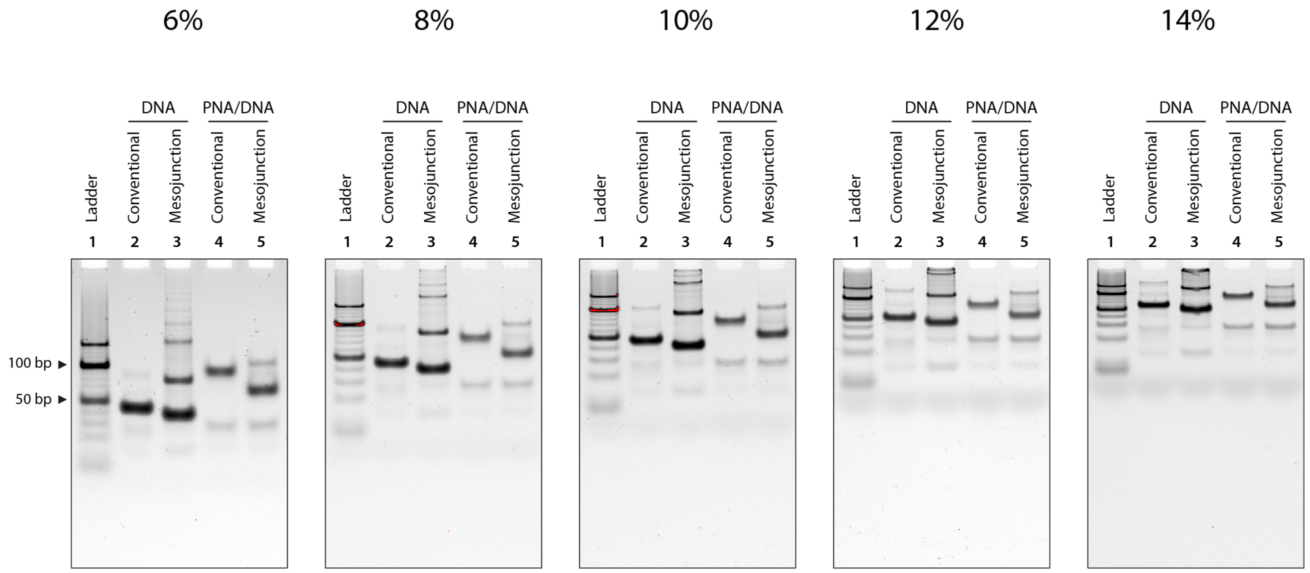


**Figure S4**. Gels containing different percentages of polyacrylamide for Ferguson analysis containing DNA and PNA/DNA junctions.

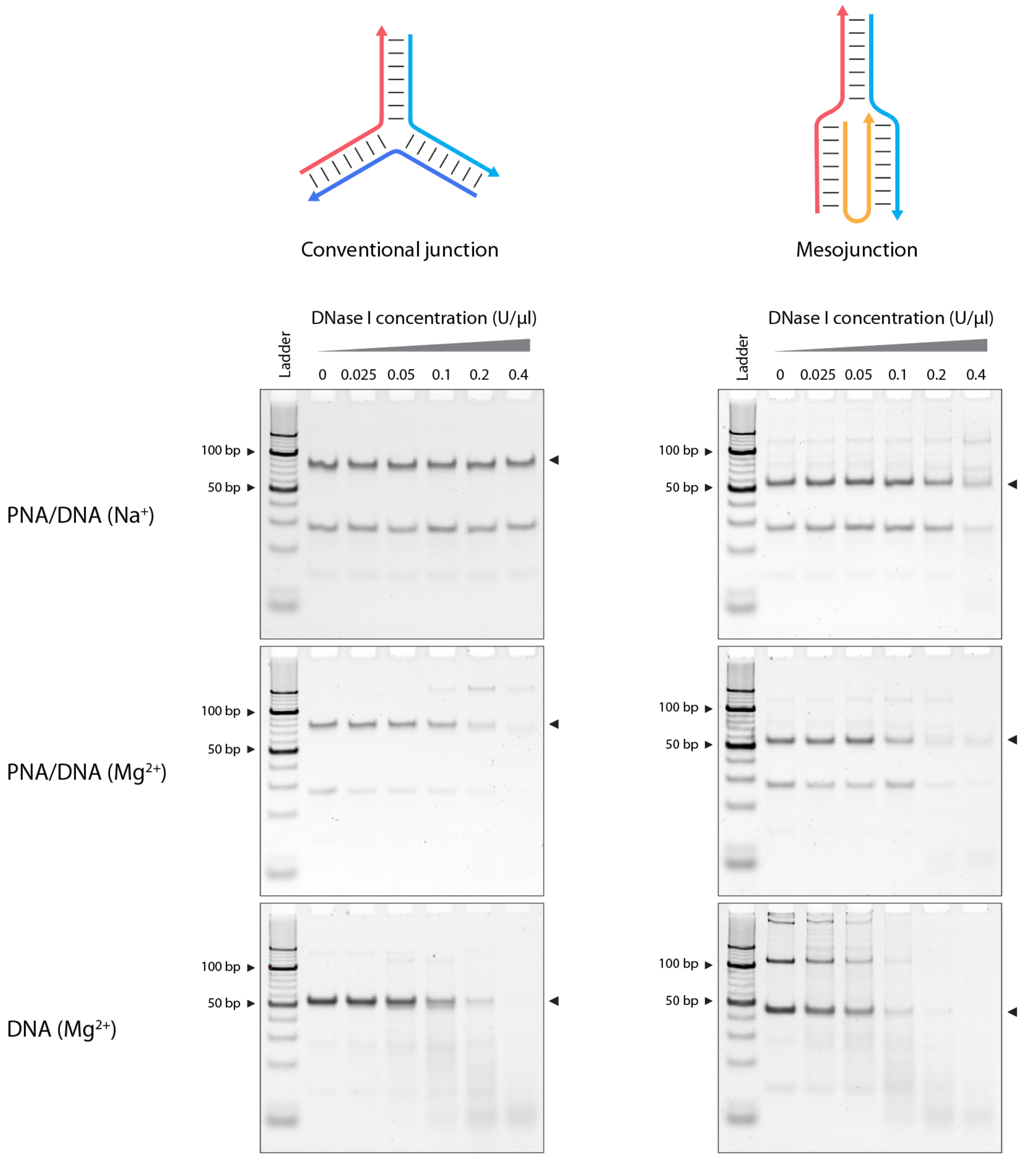


**Figure S5**. Analysis of DNase I treated DNA and PNA/DNA junctions assembled in $Mg^{2+}$ or $Na^{+}$. Full images of gels shown in Figure 4.

**Table S1**. Sequences of DNA and PNA strands used in the study.

| **Strand Name** | **Sequence (5' to 3')** |
|---|---|
| **DNA** | |
| d1-12bp | GTGCTCCGTAGCGATGGTGCGTGA |
| d2-12bp | TCACGCACCATCCTGCTACGACTC |
| d3-12bp | GAGTCGTAGCAGGCTACGGAGCAC |
| d4-12bp | GCTACGGAGCACGAGTCGTAGCAG |
| d1 | AAGTGCTCCGTAGCGATGGTGCGTGATT |
| d2 | AATCACGCACCATCCTGCTACGACTCAA |
| d3 | TTGAGTCGTAGCAGGCTACGGAGCACTT |
| d4 | GCTACGGAGCACTTTTGAGTCGTAGCAG |
| d1-16bp | GGAAGTGCTCCGTAGCGATGGTGCGTGATTGC |
| d2-16bp | GCAATCACGCACCATCCTGCTACGACTCAACG |
| d3-16bp | CGTTGAGTCGTAGCAGGCTACGGAGCACTTCC |
| d4-16bp | GCTACGGAGCACTTCCCGTTGAGTCGTAGCAG |
| **PNA (N-terminus to C-terminus)** | |
| p3 | TTGAGTCGTAGCAGGCTACGGAGCACTT-Lys-Lys |
| p4 | GCTACGGAGCACTTTTGAGTCGTAGCAG-Lys-Lys |

**Table S2**. Ferguson plot analysis.

| **Junction type** | **Slope** | **Y-intercept** |
|---|---|---|
| DNA conventional junction | -0.07321 | 0.06158 |
| PNA/DNA conventional junction | -0.07183 | -0.08355 |
| DNA mesojunction | -0.06969 | 0.05937 |
| PNA/DNA mesojunction | -0.06615 | -0.0414 |

**Table S3**. UV-melting temperatures of DNA and PNA/DNA junctions

| **Junction type** | **Tm (°C)** |
|---|---|
| DNA conventional junction | 67.8 |
| PNA/DNA conventional junction | 76.0 |
| DNA mesojunction | 64.0 |
| PNA/DNA mesojunction | 72.9 |
